# Mechanical Equilibrium of Step Transition Governs Vertical Ground Reaction Force Morphology in Human Walking

**DOI:** 10.1101/2024.07.16.603707

**Authors:** Seyed-Saleh Hosseini-Yazdi, John EA Bertram

**Affiliations:** Department of Biomedical Engineering, Schulich School of Engineering, University of Calgary, AB, Canada; Human Performance Laboratory, Faculty of Kinesiology, University of Calgary, AB, Canada; Cumming School of Medicine, University of Calgary, AB, Canada

**Keywords:** Vertical ground reaction force, Push-off, Collision, walking speed, inverted pendulum model

## Abstract

Understanding the vertical ground reaction force (vGRF) profile offers important insight into how humans regulate mechanical work during walking. Although the characteristic double-hump vGRF pattern is well documented, the mechanical factors underlying asymmetry in peak amplitudes and midstance trough timing remain unclear. Using a simple powered walking model and an inverted pendulum simulation with constant hip torque, we examined how step-transition work—collision and push-off—shapes the vGRF trajectory. We further compared these predictions to empirical data spanning walking speeds from 0.8–1.4 m. s^−1^. The simple walking model predicted symmetric vGRF profiles across speeds because collision and push-off impulses were equal, resulting in passive single-support motion. In contrast, adding hip torque within the pendular model produced stance-phase asymmetries, shifting the vGRF trough earlier when torque added energy and later when torque dissipated energy. Empirical analysis revealed that collision and push-off impulses were generally unequal except at one speed, producing asymmetric vGRF peaks. At low speeds, push-off exceeded collision; at high speeds, the reverse occurred, consistent with a need for compensatory single-support positive work. These mechanical imbalances predicted systematic shifts in trough timing toward the dominant impulse. Therefore, we propose the Vertical GRF Trough Timing Index (vGRF-TTI), combined with collision and push-off peak amplitudes, as a clinically meaningful outcome capturing the balance of step-transition work. Earlier troughs with elevated collision peaks indicate impaired push-off or constrained gait conditions, whereas later troughs with larger push-off peaks reflect compensatory or enhanced propulsion. These metrics provide sensitive, mechanism-based indicators of gait efficiency and neuromotor control.

## Introduction

Walking is performed by exerting forces on the terrain. The reaction of the induced forces (Ground Reaction Forces: GRF) accelerates the walkers’ Center of Mass (COM) in the direction of progression (Muller 2018). The vertical direction component is the largest since the legs support body weight (Masani, Kouzaki, and Fukunaga 2002), as a result, the analysis of vertical GRF has attracted much attention (Libera, Streamer, and Queen 2025; Schwartz, Rozumalski, and Trost 2008). Triaxial GRF can be collected on instrumented treadmills (Hosseini-Yazdi and Kuo 2024) in the lab setting but requires considerable upfront capital expenditure. Thus, many researchers may utilize inexpensive uniaxial force plates (Voloshina et al. 2013) or shoe insoles (Libera, Streamer, and Queen 2025) that only record vertical force to avoid lab space and cost limitations. Vertical GRF profiles are affected by several factors, such as terrain complexity (Aminiaghdam and Rode 2017; Hosseini-Yazdi and Kuo 2025), view of approaching substrate irregularities (Hosseini-Yazdi Seyed-Saleh 2024; Majlesi, Farahpour, and Robertson 2020), age (Laufer 2005), etc. However, a limited mechanical explanation has been provided for those circumstances when the two vertical GRF peak amplitudes are different and the midstance trough is skewed, i.e. when the vertical GRF profile is asymmetric (Figure 1).

**Figure 1:**
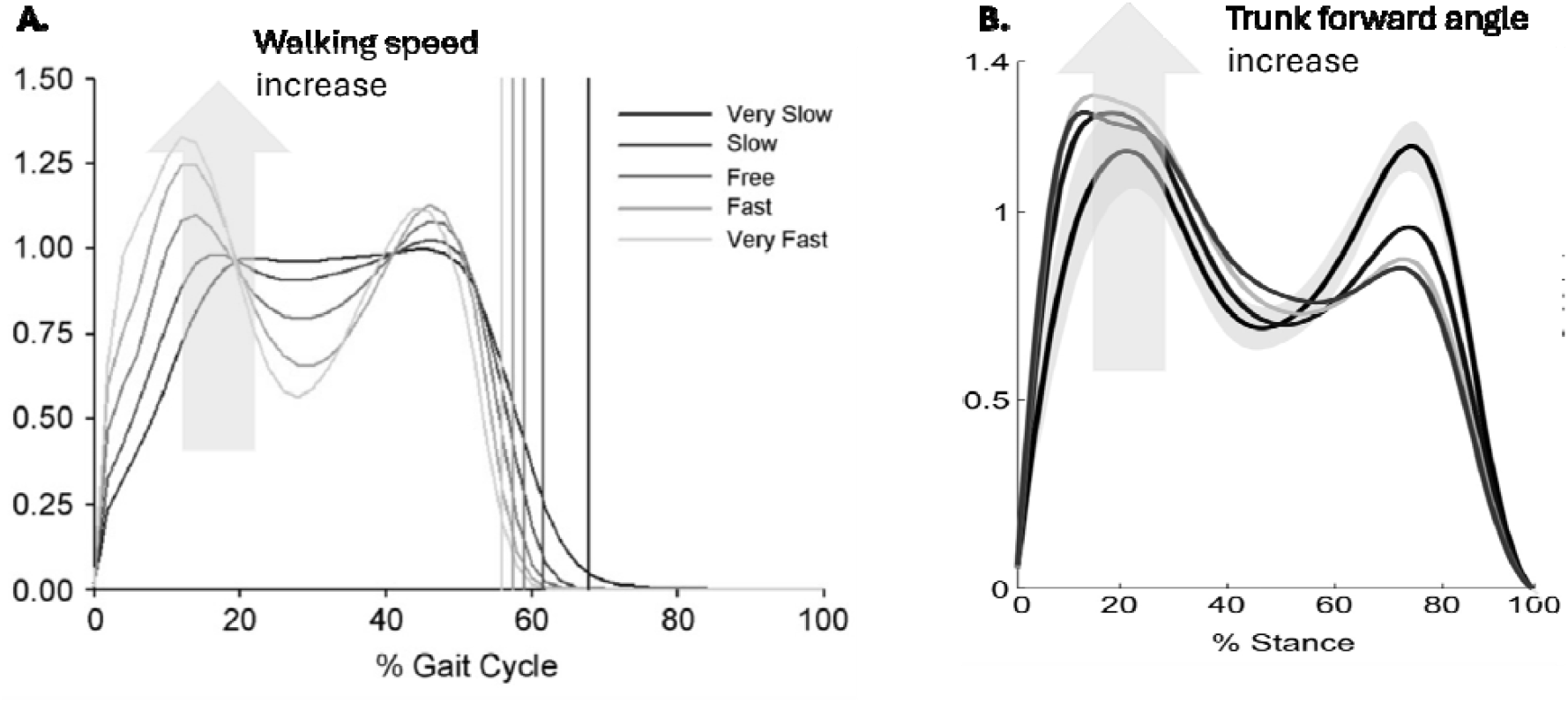
Vertical ground reaction force (GRF) of walking changes when walking conditions are altered: (A) when walking spee is varied (adopted from (Schwartz, Rozumalski, and Trost 2008)), (B) when the truck forward angle is raised (adopted from (Aminiaghdam et al. 2017)). The solid black line represents free normal walking. While the vertical GRF for nominal walkin demonstrates a fairly symmetrical profile, deviations lead to asymmetrical trajectories.

It is suggested that the first peak of the vertical ground reaction force is due to weight acceptance (Aminiaghdam et al. 2017; Libera, Streamer, and Queen 2025), i.e. a collision as the heel strike of the next stance limb affects the COM velocity vector (Adamczyk and Kuo 2009), and the second is due to ankle-generated force to accelerate the COM forward and upward (Adamczyk and Kuo 2009; Muller 2018). Various factors may affect the magnitude of vertical GRF peaks. For example, carrying extra load distorts the vertical GRF profile (Aminiaghdam and Rode 2017; Huang and Kuo 2014). Aminiaghdam et al. (Aminiaghdam and Rode 2017) have suggested that greater loads and loading rates or lower unloading rates alter the vertical GRF peaks. It is also proposed that walking with a crouched posture leads to higher vertical GRFs and loading rates during early stance and lower vertical GRFs during the pre-swing phase (Aminiaghdam et al. 2017). It is further suggested that, modeling the human leg as a spring and damper, the damper shifts the vertical GRF later in stance by increasing forces after the heel strike and decreasing forces at toe-off, resulting in an early lift-off (Aminiaghdam et al. 2017). One factor that is sometimes often not appreciated properly is that the vertical GRF must equal (or be very close to) body weight over the complete stride – whatever speed the individual is walking, it is necessary to support their weight against gravity while not producing excess force that would accelerate their COM beyond foot contact.

In addition to load support, GRF represents the work performed by the legs on the COM to sustain progression. It has been shown that human walking involves a step-to-step transition during double stance over which the direction of COM velocity changes (Adamczyk and Kuo 2009). This velocity change requires energy dissipation (Kuo 2002). Therefore, if the kinetic energy is to be preserved, the lost energy must be compensated for. Such energy dissipation and restoration are observed during the step transition (Donelan, Kram, and Kuo 2002a, 2002b) by force application of leading and trailing legs generating the characteristic double hump shape (Adamczyk and Kuo 2009). The two limbs work in concert to manage the transition where the toe-off (push-off) force peak of the trailing leg is followed by the leading leg’s heel-strike peak. This sequence (push-ff preceding heel strike) is important because it minimizes step transition energy dissipation (Kuo 2002), therefor walking energetic cost (Donelan, Kram, and Kuo 2002a) (Figure 2).

**Figure 2:**
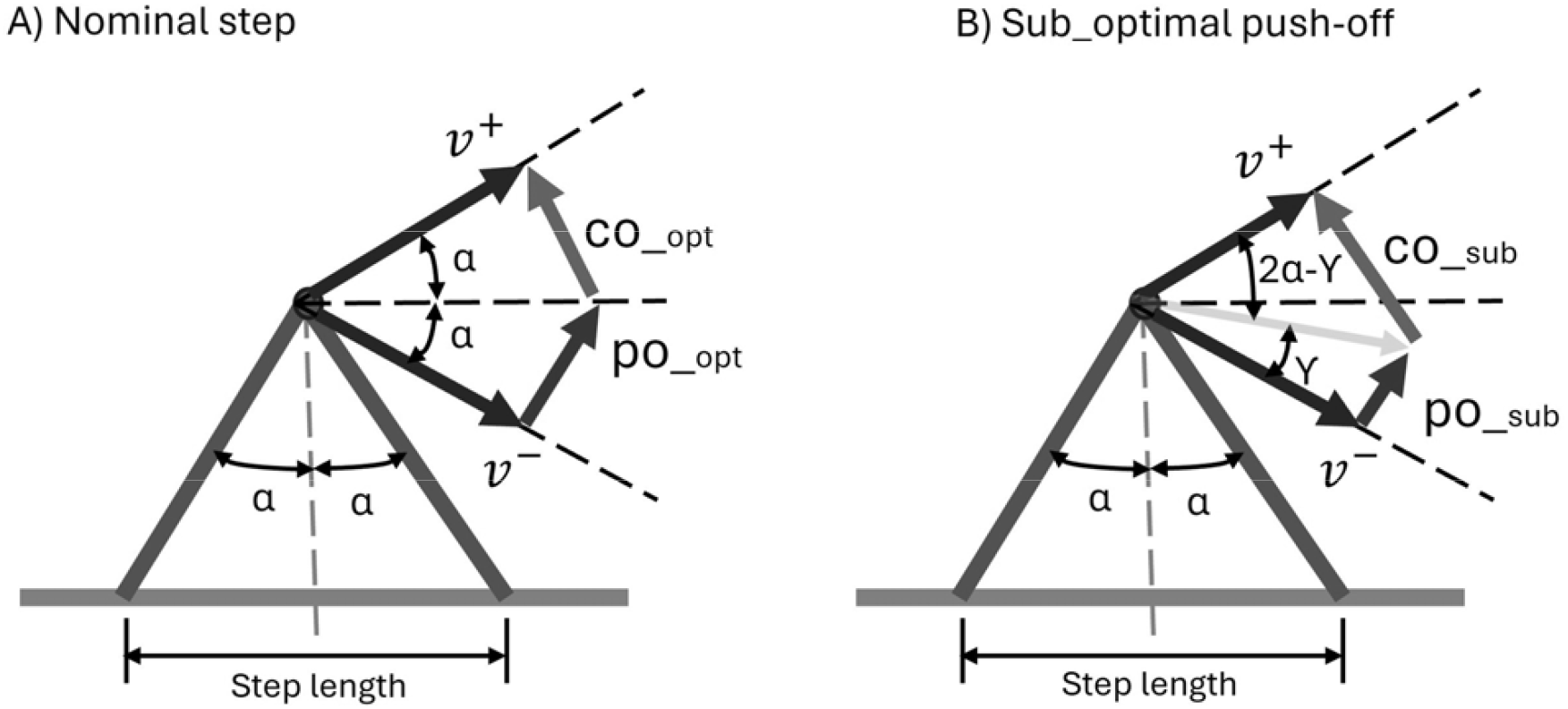
A simple model of powered walking that explain the step-to-step transition required work performance. (A) in nominal walking, the energy compensation and energy dissipation are equal. (B) An example of deviation from nominal walking is when for a given speed, the magnitude of pre-emptive push-off that compensates step transition dissipation is less than required.

Vertical GRF peak amplitude asymmetry has not been fully explained, especially where the factors involved are verified by experimental data. We hypothesize that the amplitude of vertical GRF peaks at heel-strike and toe-off are determined by the walking speed collision and push-off magnitudes. Hence, when walking speed deviates from the mechanically optimal (Hosseini-Yazdi and Bertram 2025a), the two vertical GRF peak amplitudes become unequal. Due to its relative magnitude, understanding the vertical GRF profile may provide substantial insight into human locomotion (Fineberg et al. 2013). In this study, we first attempt to understand its variation using a simple walking and pendular model for different walking conditions. Further, we compare the simulation results with analysis of empirical data available in the literature.

## Materials and Methods

Since the vertical GRF magnitude in walking is relatively larger than other force components, the step-to-step transition work is often associated with its application (Adamczyk and Kuo 2009)(Figure 3). As a result, we focus on its role using a simple walking model simulation (Kuo 2002) and a pendular model (R. McN. Alexander 1995; R.McNeill Alexander 1992). We then compare the model results to empirically measured results (Hosseini-Yazdi Seyed-Saleh 2024).

**Figure 3:**
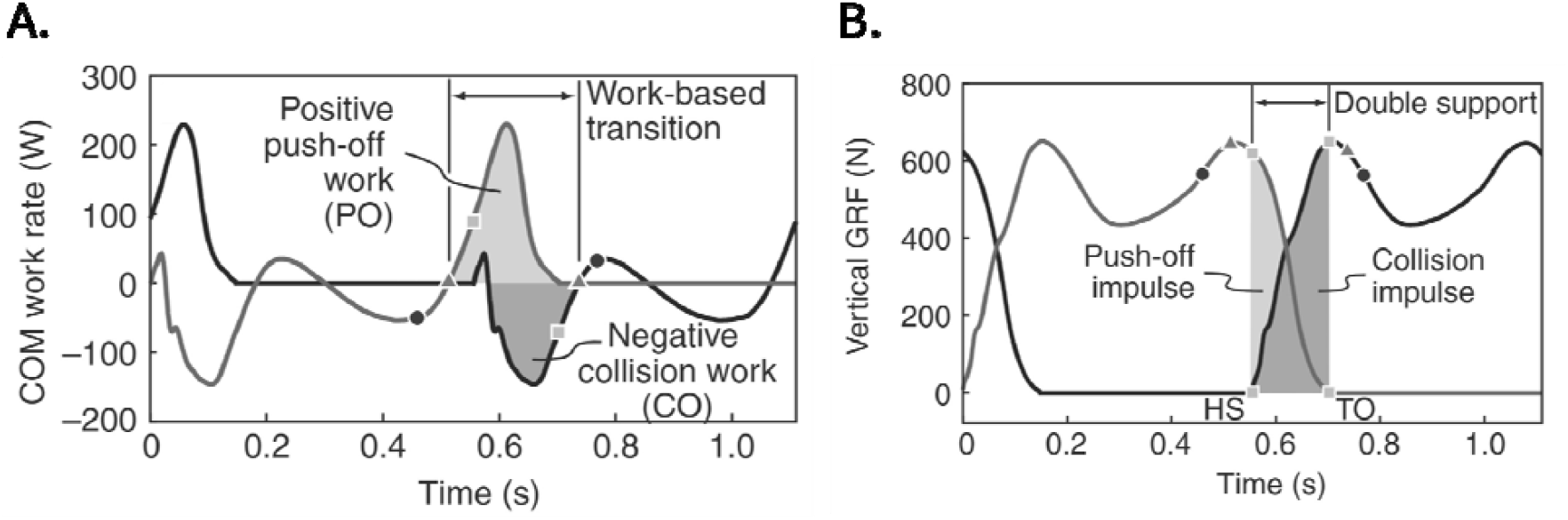
Walking step-to-step transition involves a simultaneous positive and negative work performance to adjust the COM velocity for the new stance while minimizing required mechanical work and as such metabolic cost of walking. (A) step transition mechanical work by leading and trailing legs, (B) step transition impulses to perform step transition (adopted from (Adamczyk and Kuo 2009).

### Simulation

Our first analysis uses a simple powered walking model that assumes concentrated mass in the pelvis with rigid massless legs (Kuo 2002). It assumes that the entire step work, positive and negative, occurs during the step transition when the previous stance leg push-off is followed by the collision of the next stance leg. For a given walking speed (*v*_*ave*_), the push-off impulse is *po* = *v*_*ave*_. tan *α* in which ‘*α*’ represents half the angle between leading and trailing legs at the point of transition (Figure 2).

We break down the stance phase into three sub-phases; first double support representing the heel-strike collision impulse, single support with no active work as an inverted pendulum, and the second double support demonstrating the toe-off push-off impulse. Therefore, transition impulses (co: for heel-strike or collision and po: for toe-off or push-off) can be represented as the area under the force trajectory of the associated region. Thus, the step transition impulses can be proxies for the amplitude of collision and push-off peaks, since the model assumes a very short duration for impulse application (Kuo 2002) (Figure 4). We assume that the collision ends at the first vertical GRF whereas push-off starts at the second peak (Adamczyk and Kuo 2009) (Figure 3).

**Figure 4:**
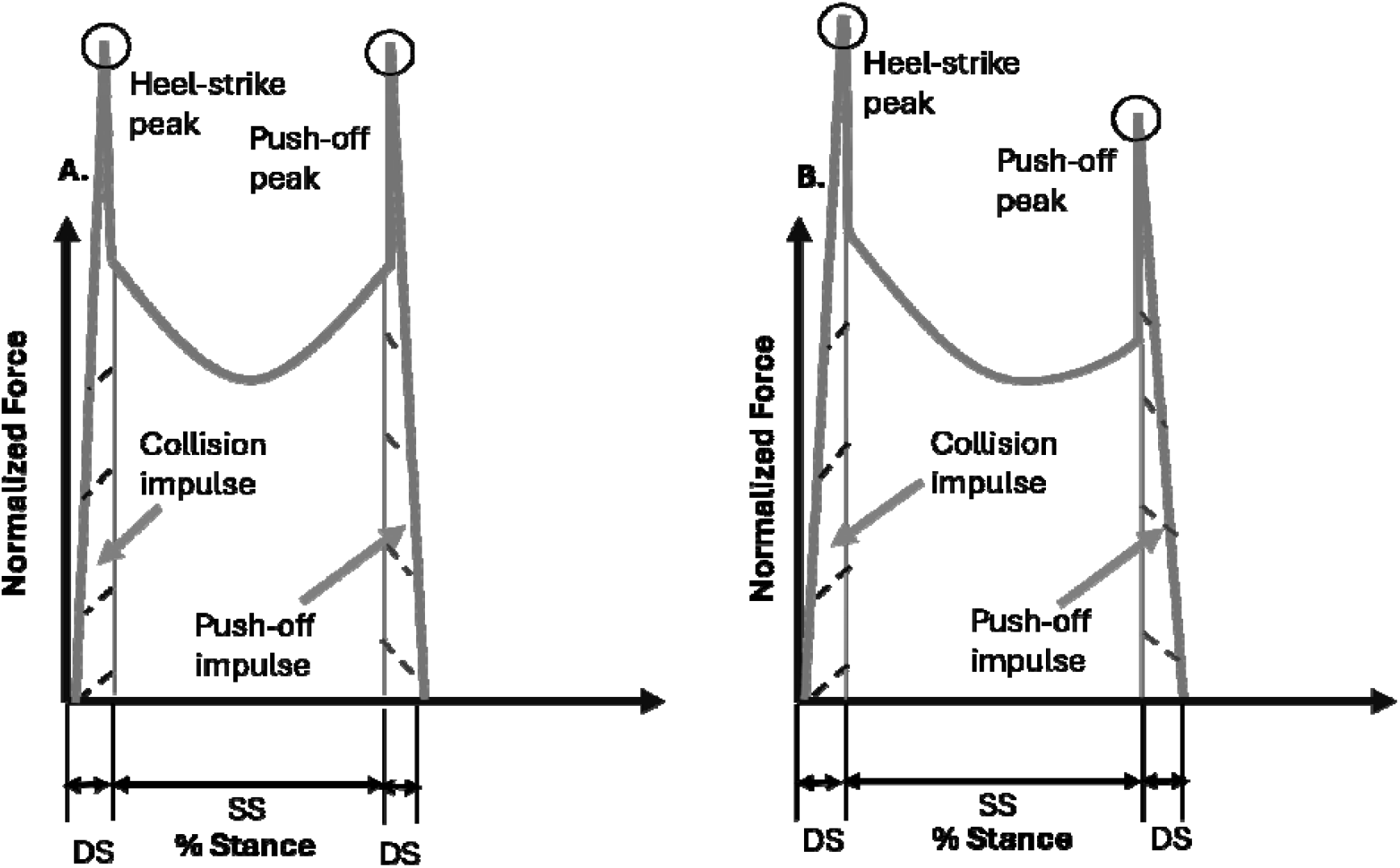
A vertical GRF during a walking stance may be described by heel-strike weight acceptance collision, single support motion like an inverted pendulum, and the push-off. According to the model, the durations for active force applications are very short, therefore, for the sake of representation vertical force trajectories depicting active impulses are plotted close to vertical direction. (A) when vertical GRF is symmetrical, (B) when vertical GRF is asymmetrical.

During the single support phase, the motion of the COM is proposed to closely resemble an inverted pendulum (Cavagna, Heglund, and Taylor 1977; Kuo, Donelan, and Ruina 2005a). Having short double support durations (R. McN. Alexander 1995; Kuo 2002), it is fair to consider pendular motion for the entire stance (Figure 5). As such, the pendulum normalized axial force (*T*) can be represented as (Renjewski 2025):

**Figure 5:**
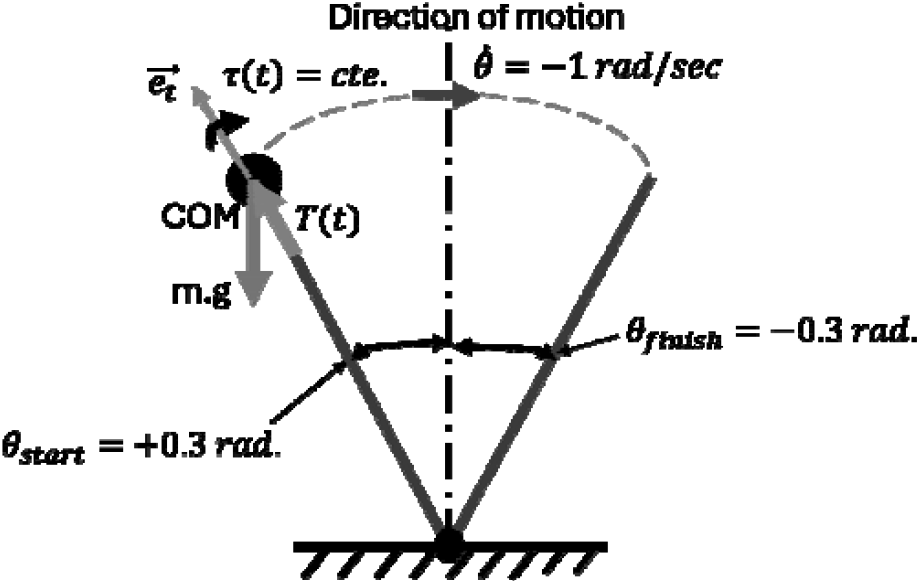
the simulation of single support with inverted pendulum motion in which all the mass is concentrated and legs are rigid. A constant torque is applied to the COM to modulate its motion. It is to represent the hip work.

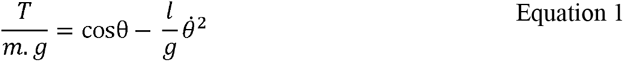

Therefore, the vertical component of the reaction forces (*F*_*Z*_) is:

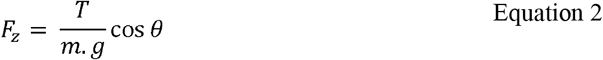

For the sake of this analysis, we assume that the pendular motion starts at *θ* = +0.3 *rad*., ends at *θ* = −0.3 *rad*., and the angular velocity is 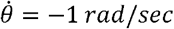. As a small increase in the complexity of the model, we assume that single support active work is exerted by a constant hip torque (Hosseini-Yazdi and Bertram 2025b; Kuo, Donelan, and Ruina 2005a; Voloshina et al. 2020). We consider three hip torque magnitudes (*τ*(*t*)∼ *cte*.): -0.15, 0, and 0.15 to evaluate the effect of single support work on the profile of the vertical GRF. Therefore, the equation of motion for the inverted pendulum becomes (Kuo 2002):

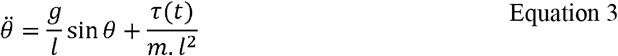

Since, the COM motion based on the right-hand rule is in the negative direction, the positive torque reduces the energy of the pendulum while negative torque increases it. We also check if the vertical reaction force is symmetrical or is skewed to a particular direction using a Parabola Skewness Index: (PSI) where if the vertex is shifted to the right PSI > 0, if the vertex is shifted to the left PSI < 0, for complete symmetry PSI is zero (|PSI| < 0.001). Here, the ‘*l*’ stands for the leg length (here 1 *m*) whereas the ‘*g*’ represents gravitational acceleration.

Since this simulation can provide quantification of the GRF asymmetry, they may serve as a basis for developing a simple metric to indicate the underlying mechanisms responsible. One objective of this analysis is to devise such a simple metric for the purpose of applying to GRF profile analysis.

### Empirical Data Analysis

During stance, four distinct phases in the COM power profile can be recognized (Kuo, Donelan, and Ruina 2005a). While collision and push-off are the negative and positive work during double support, rebound and preload are the positive and negative work during single support (Hosseini-Yazdi and Bertram 2025a) (Figure 6). We consider collision and push-off as the result of associated impulses (Adamczyk and Kuo 2009; Kuo 2002). Similar to the simulation, we assume that the collision ends at the first vertical GRF while push-off starts at the second peak (Adamczyk and Kuo 2009) (Figure 3 and Figure 4). Therefore, collision and push-off impulse magnitudes define the vertical GRF first and second peak amplitudes.

**Figure 6:**
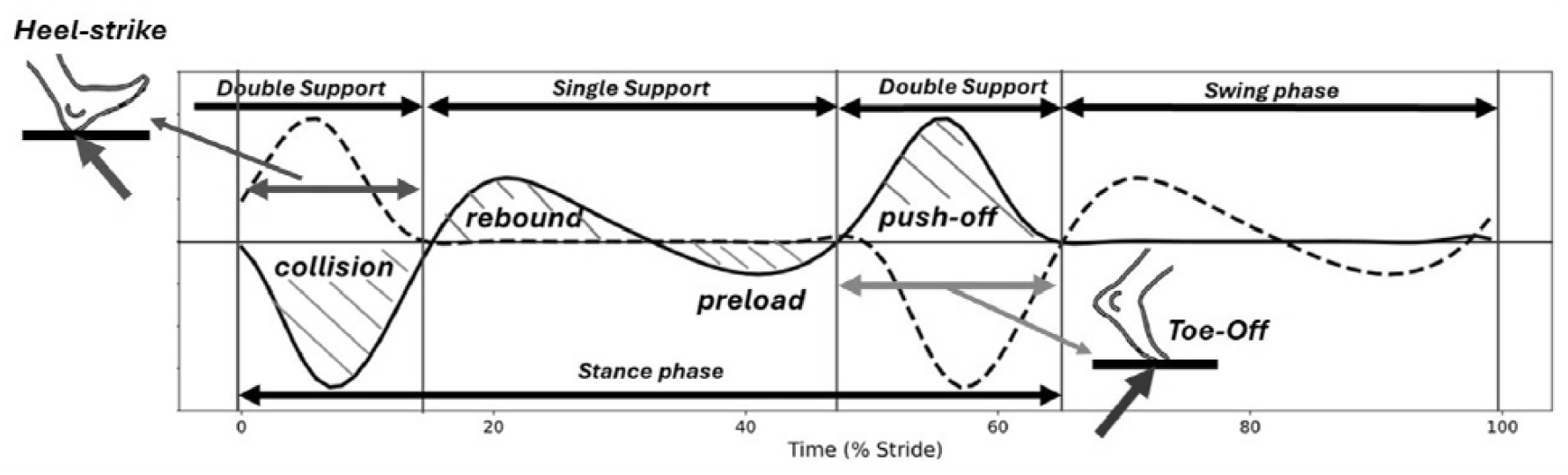
During stance phase, the COM power depicts four distinct sub-phases. It starts with a collision of the heel-strike durin the first double support. It is followed by the positive and negative work of single support, i.e. rebound and preload. It ends with the push-off that infuses positive work to COM (adopted from *(Hosseini-Yazdi and Bertram 2025a)*)

We utilize available empirical data suggesting trajectories for collision and push-off when walking speed varies (0.8 m. s^−1^ to 1.4 m. s^−1^) (Hosseini-Yazdi Seyed-Saleh 2024). We remove the random effect of treadmill calibration (Hosseini-Yazdi and Bertram 2025b, 2025a). For any speed, the step transition impulses are calculated by 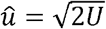, where ‘U’ and ‘û’ are work and its associated impulse, respectively (Darici, Temeltas, and Kuo 2018).

## Results

Based on the simulation, we observed that with walking speed increase (0.8 m. s^−1^ to 1.4 m. s^−1^), the step transition impulses grew from 0.20 m. s^−1^ to 0.57 m. s^−1^. The step transition collision and push-off also rose from 0.02 J. kg^−1^ to 0.16 J. kg^−1^. As such, the simulation demonstrated that the mechanical energy dissipation of collision and the active work of push-off were equal for any speed in this range (Figure 7). Thus, the single support after the step-transition did not require any regulatory active work. In other words, the simulation predicted symmetrical profiles for walking GRF.

**Figure 7:**
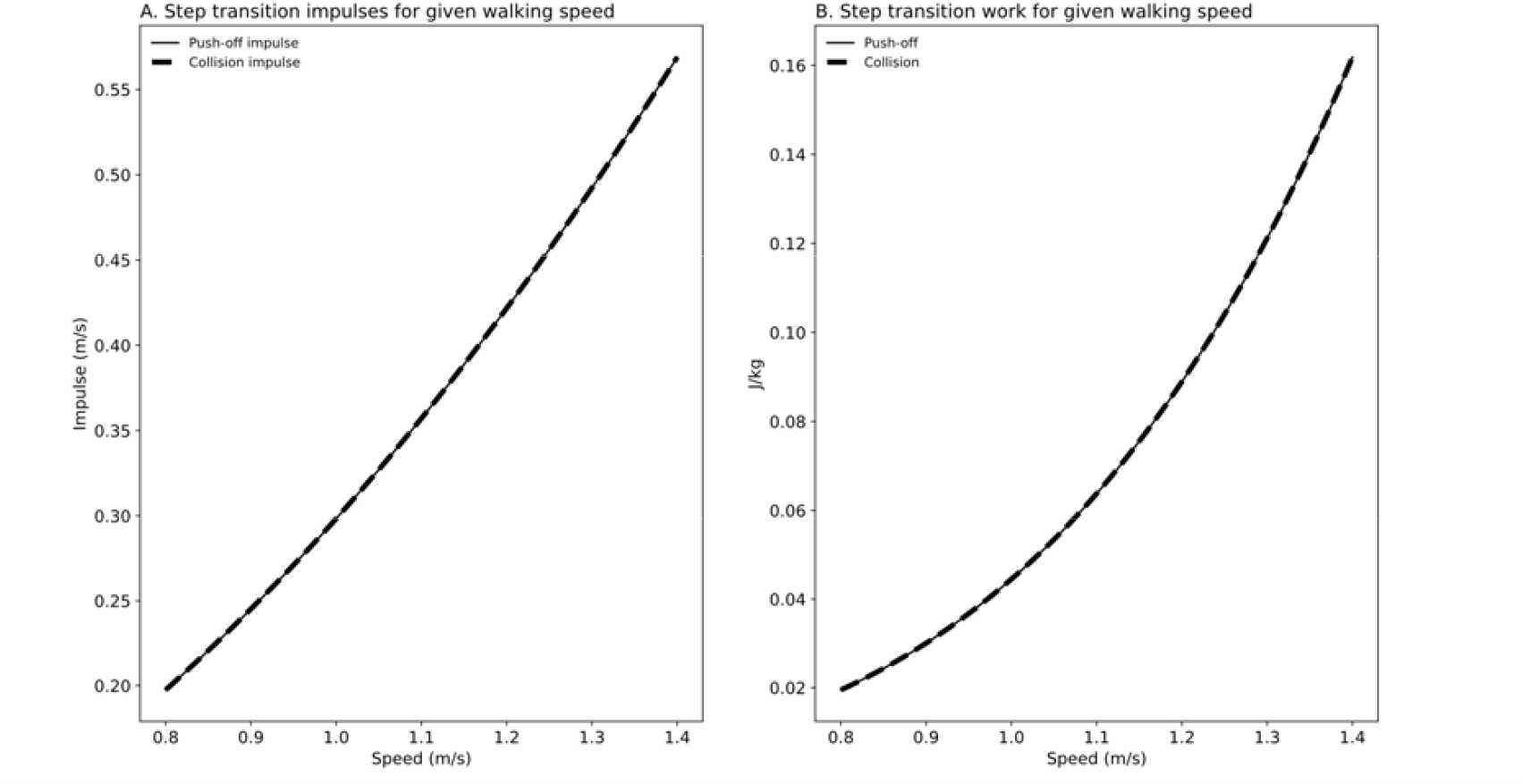
Simple walking model simulation prediction for (A) the step transition impulses, (B) step transition work for given walking speed range. For any speed, the model assumes equal dissipation and energy compensation to maintain the walker’s mechanical energy during the step transition, as such the single support may be done passively. Therefore, it predicts a symmetrical vertical GRF profile for any given speed.

The simulation shows similar profiles for total pendular axial force and vertical reaction force. Without hip torque, the duration of stance was 1.09 sec while normalized pendular axial force and vertical reaction were 0.99 and 0.98, respectively. With a hip torque that resisted the pendular motion (positive direction), the stance duration increased to 1.51 sec (+%38.5) while the changes in axial force (*T* = 1.00, or +%1.0), and vertical reaction (*F*_*Z*_ =0.99, or +%1.0) were relatively small. On the other hand, with a hip torque enhancing motion (in the negative direction), the stance duration declined to 0.93 sec (-%17.2) while the axial force (*T* = 0.98, or - %1.0) and vertical reaction (*F*_*Z*_ =0.97, or -%1.0) reduced slightly. We also observed asymmetries when pendular motions were performed along with hip torque. When there was no hip torque, the start and end vertical reaction forces were 0.82 with PSI of 0.000 while with supporting (in the negative direction) and opposing (in the positive direction) torques the ending vertical reaction forces were 0.79 (PSI = -0.013, skewness to left or earlier stance) and 0.83 (PSI = 0.012, skewness to right or later stance), respectively. In other words, supporting hip torque, skewed the pendular force peak toward early stance, whereas an opposing torque pushed it towards the later stance. This further represented asymmetries when a hip torque was applied during pendular motion (Figure 8).

**Figure 8:**
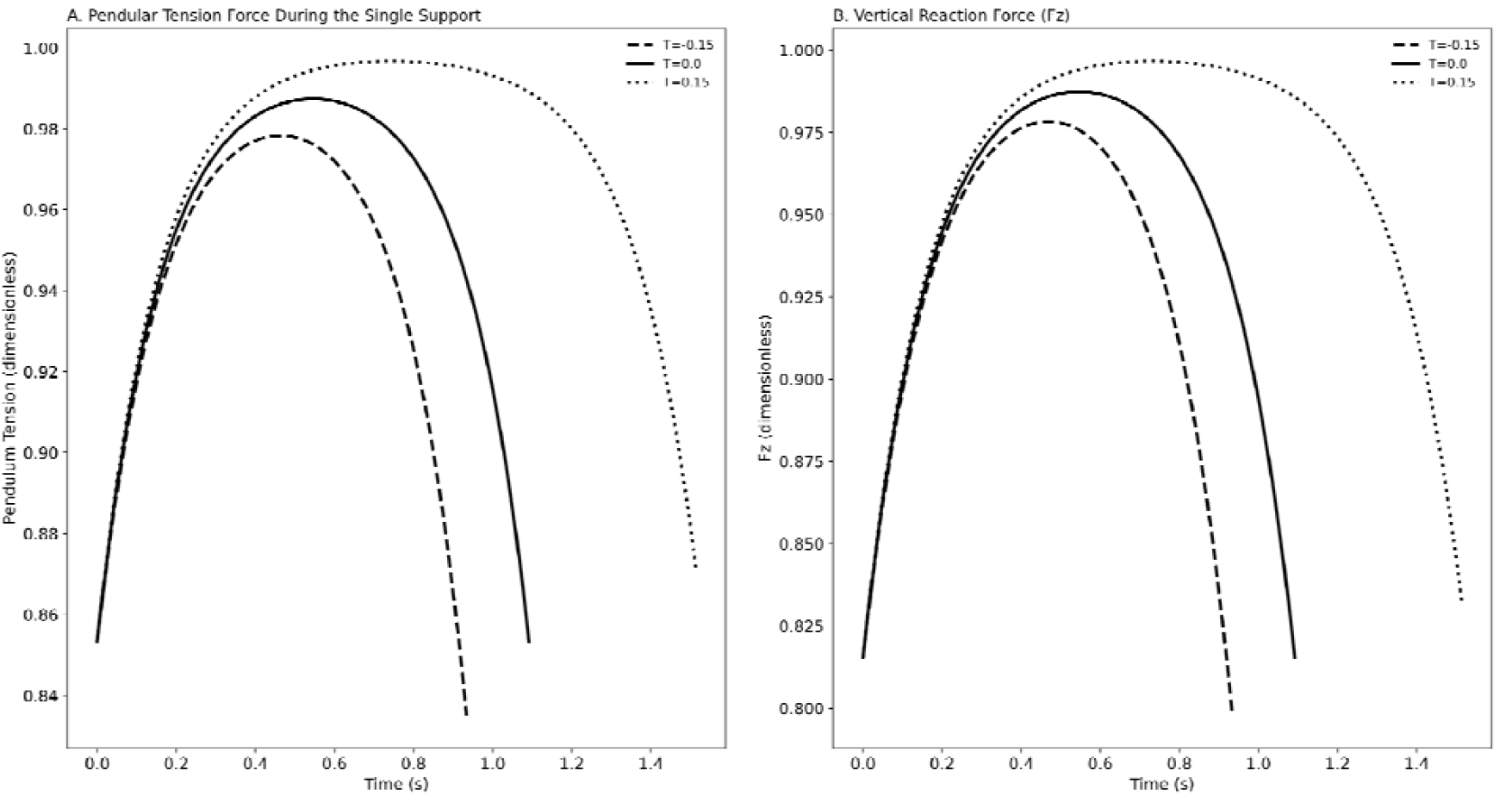
The simulation of single support motion with an inverted pendulum when hip torque is present. The simulation suggests that the presence of hip torque changes the magnitude of peak force and trajectory, yet the influence on the duration of stance, i.e. force profile is more pronounced. The hip torque application also affects the symmetry of force profile skewing it based on the direction of torque. (A) Total pendulum axial force, (B) vertical reaction force of pendulum.

In contrast to the simple walking model, the empirical data analysis predicted asymmetrical GRF peaks except for one particular walking speed. As walking speed increased, the associated push-off and collision varied from 0.16 J. kg^−1^ to 0.38 J. kg^−1^, while the collision dissipation magnitude varied from 0.06 J. kg^−1^ to 0.44 J. kg^−1^. As a result, the associated heel-strike and toe-off impulses ranged from 0.41 m. s^−1^ to 0.61 m. s^−1^, and from 0.24 m. s^−1^ to 0.67 m. s^−1^, respectively (Figure 9). Therefore, at lower walking speeds, the push-off peaks were greater than those of the heel-strike collisions, whereas at elevated speeds the order was reversed. Therefore, single support required regulating active work. At *v* = 1.21 m. s^−1^, the collision and push-off impulses were equal, and thus single support did not require active work. In other words, the heel-strike and push-off peaks were dissimilar except at the designated speed where the magnitudes of collision and push-off were equal.

**Figure 9:**
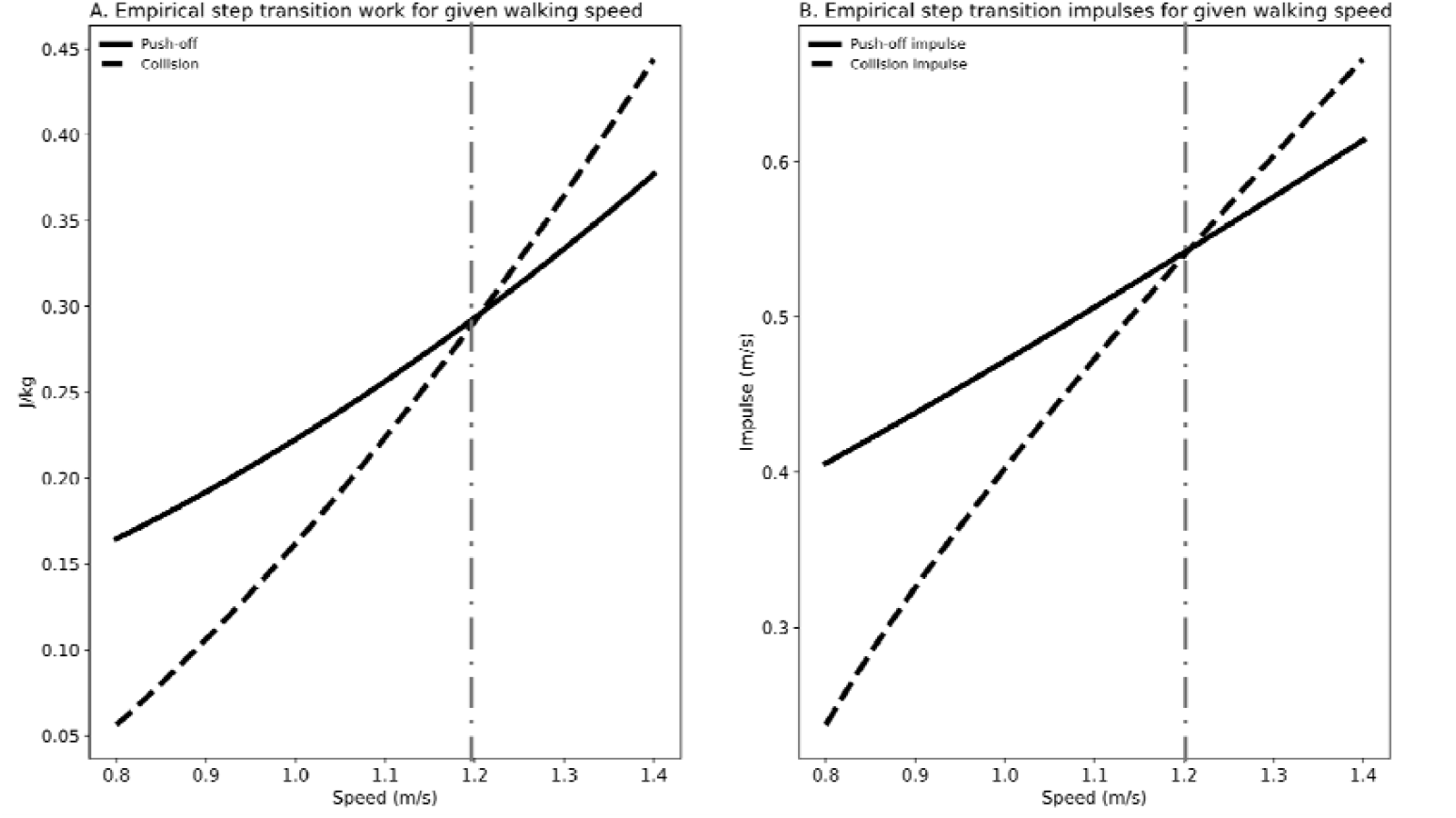
The analysis of empirical data for step transition (A) work, (B) derived impulses. The analysis suggests that there is only one walking speed (designated at) for which the magnitudes of energy dissipation and compensation are equal. As such, the associated vertical GRF must have a symmetrical profile with inverted pendulum like motion for the single support phase in which no active muscle work is performed. Therefore, for speeds less than the designated value, the push-off peak is predicted to be larger than collision peak. The order is predicted to be reverse for speeds higher than the designated value.

## Discussion

GRF analysis is the simplest approach to study human walking (Park and Kim 2022) and is utilized for clinical biomechanic evaluations (Campanelli 2023). GRF indicates the instantaneous acceleration of the COM. The GRF profiles are affected by walking conditions since humans modulate their walking by exerting forces and performing mechanical work (Kuo, Donelan, and Ruina 2005b). Since vertical GRF is the dominant force during human walking (Libera, Streamer, and Queen 2025), we have attempted to provide a mechanistic explanation of its trajectory as walking conditions are altered.

In this study we have used a simple walking model focusing on the step-to-step transition (Kuo 2002). Then, we utilized a pendular model (R. McN. Alexander 1995; Cavagna, Heglund, and Taylor 1977) with active torque applied at the hip. Although the pendular model is able to predict the horizontal force of walking fairly reliably (Buczek et al. 2006), it does not show the “M” shape of the vertical force component seen in human walking. The pendular model presents a concavity that is opposite of what is observed experimentally (Campanelli 2023). However, it provides valuable insights about human walking. The pendular model suggests that the single support phase is performed with minimal torque, i.e. free or passive motion (R.McNeill Alexander 1992; Cavagna, Heglund, and Taylor 1977; Kuo, Donelan, and Ruina 2005a). Nevertheless, there are studies that highlight the role of single support work to regulate COM mechanical energy throughout stance (Hosseini-Yazdi and Bertram 2025a, 2025b, 2025c). Therefore, we have evaluated the impact of constant torque (Voloshina et al. 2020) for a pendular model, whether opposing or supporting the COM motion. Finally, we utilized available empirical data to predict the symmetry or asymmetry of vertical GRF based on the step transition work and subsequent mechanical energy imbalance.

The simulation based on the simplest walking model (Kuo 2002) suggests a symmetrical force profile for any adopted walking speed. Since this model assumes that the entire step active work occurs during a short duration at the step-to-step transition (Donelan, Kram, and Kuo 2002a). The simplest walking model assumes completely passive motion over the single support phase (Cavagna, Heglund, and Taylor 1977). Thus, this model suggests nominal walking for any adopted speed. Since it is shown that active work is not limited to the step-transition only (Hosseini-Yazdi and Bertram 2025a, 2025c), we may consider adding work through the application of torque at the hip as a modified approach to enhance the simple walking model capabilities (Darici and Kuo 2022; Darici, Temeltas, and Kuo 2018, 2020).

Constant hip torque leads to changes in vertical reaction force as work is regulated over single support. We observe that torque application mainly affects the time duration of single support and, as such, the magnitude of applied impulses. The amplitudes of peak forces also change, and the force profiles become asymmetric. Based on this simulation, we expect the vertical GRF trough to shift to later stance when the applied torque dissipates COM energy, whereas it shifts the GRF trough toward early stance when hip torque adds to COM energy.

Based on empirical data analysis, for a given walking speed, we observe an imbalance in step-transition work. At elevated walking speeds when the collision exceeds the push-off, there must be a mechanical energy addition during the single support phase (Hosseini-Yazdi and Bertram 2025a). Since with walking speed increase, the duration of double support declines (Williams and Martin 2019), the step-transition work must occur in a shorter time frame. As such, we expect dominant heel-strike peaks compared to push-off. We also expect that the vertical GRF trough to be skewed towards the collision peak. This prediction is supported by examining associated vertical GRF profiles (Nilsson and Thorstensson 1989; Schwartz, Rozumalski, and Trost 2008).

On the contrary, when the push-off is greater than the collision at reduced walking speeds, there must be energy dissipation during single support. Hence, we expect a dominant push-off peak versus collision with the single support GRF trough skewed toward push-off. Although experimental data supports the skewness of the trough toward later stance (Nilsson and Thorstensson 1989), it is hard to recognize a dominant push-off peak (Schwartz, Rozumalski, and Trost 2008). The extension of the double support phase with reduced speed (Williams and Martin 2019) may spread push-off impulse application over an expanded time frame. Therefore, the push-off peak amplitude is reduced.

When step transition work is balanced, i.e. collision = push-off, the vertical GRF must be symmetrical. As such, not only are the amplitudes of push-off and collision peaks equal, but also the single support phase must closely resemble an inverted pendulum with passive motion (zero hip torque) (R. McN. Alexander 1995). Therefore, the single support force must be symmetrical with the trough at midstance. It is suggested that the self-selected walking speed coincides with the mechanically preferred walking speed (Hosseini-Yazdi and Bertram 2025a) during which step-transition work is balanced, and there is no active work during single support (Hosseini-Yazdi and Bertram 2025b). The vertical GRF of free walking depicts a fairly symmetrical profile (Schwartz, Rozumalski, and Trost 2008).

Humans modulate their gaits to adjust to the imposed walking conditions (Hosseini-Yazdi Seyed-Saleh 2024). One method is to adjust the force generated to perform active work. Although force application can have a large influence, not all modulation happens with force exertion. This is especially the case in short time intervals. Nonetheless, GRF analysis is a powerful tool that enables the search for other gait compensation methods or recognize possible deficits that possibly could be used for rehabilitation or gait training. The magnitude and ease of measurement (Libera, Streamer, and Queen 2025), makes vertical components the first target for gait analysis.

We suggest that the vertical GRF profile is affected by the step-transition and subsequent single support active work. Only when walking is mechanically preferred (Hosseini-Yazdi and Bertram 2025a) isthe vertical GRF symmetrical. Any walking condition that prohibits or disrupts push-off must lead to an increase in collision dissipation (Hosseini-Yazdi and Bertram 2025b). Therefore, the active work of single support shifts the vertical GRF trough closer to heel-strike (early stance). Such vertical GRF trajectories may be observed during challenging walking conditions like uneven walking (Hosseini-Yazdi and Kuo 2025), restricted lookahead or with age (Hosseini-Yazdi Seyed-Saleh 2024). On the contrary, when push-off is more than collision, the midstance trough is shifted toward toe-off (later stance).

A clinically meaningful outcome from our study is the Vertical GRF Trough Timing Index (vGRF-TTI)—the percentage of stance at which the minimum vertical GRF occurs—combined with the amplitude of the early (collision) and late (push-off) vertical GRF peaks. Together, these features capture the mechanical balance of step-to-step transition. When push-off is reduced or disrupted—such as with aging, pathology, uneven terrain, or restricted lookahead—the trough shifts earlier toward heel-strike and the collision peak increases while the push-off peak diminishes, indicating elevated collision energy loss and insufficient single-support active work. Conversely, when push-off exceeds collision, the trough shifts later toward toe-off and the push-off peak becomes larger relative to the collision peak, reflecting enhanced or compensatory late-stance power generation. Because symmetrical vertical GRF profiles only arise during mechanically preferred walking, deviations in trough timing and peak amplitudes provide sensitive, clinically accessible indicators of gait inefficiency, impaired neuromotor control, and altered mechanical work strategies.

